# Assessment of Treatment-Specific Tethering Survival Bias for the Juvenile blue crab *Callinectes Sapidus* in a Simulated Salt Marsh

**DOI:** 10.1101/2023.01.25.525559

**Authors:** Cole R. Miller, A. Challen Hyman, Daniel Shi, Romuald N. Lipcius

## Abstract

The blue crab (*Callinectes sapidus*) is ecologically and economically important in Chesapeake Bay. Nursery habitats, such as seagrass beds, disproportionately contribute individuals to the adult segment of populations. *Spartina alterniflora* salt marshes are intertidal nursery habitats which may serve as a refuge from predation for juvenile blue crabs. However, the effects of various characteristics of salt marshes on nursery metrics, such as survival, have not been quantified. Comparisons of juvenile survival between salt marshes and other habitats often employ tethering to assess survival. Although experimental bias when tethering juvenile prey is well recognized, the potential for habitat-specific bias in salt marshes has not been experimentally tested. Using mesocosm experiments, we tested if tethering in simulated salt marsh habitats produces a habitat-specific bias. Juvenile crabs were randomly tethered and un-tethered in mesocosms at varying simulated shoot densities. Tethering reduced survival, and its effect was not habitat specific, irrespective of shoot density, as evidenced by a non-significant interaction effect between tethering treatment and habitat. Thus, tethering juvenile blue crabs in salt marsh habitat did not produce treatment-specific bias relative to unvegetated habitat across a range of shoot densities and survival of tethered and untethered crabs was positively related to shoot density. These findings indicate that tethering is a useful method for assessing survival in salt marshes, as with other nursery habitats including seagrass beds, algae and unstructured sand.

## Introduction

A major objective in estuarine ecology and fisheries science is understanding the influence of habitat on the population dynamics of commercially exploited fisheries. In particular, ecologists have emphasized identifying critical juvenile habitats largely because early life stages are the most vulnerable for fish and aquatic invertebrates (Beck et al., 2001; Dahlgren et al., 2006; Nagelkerken et al., 2015). As the extent and quality of productive coastal habitats is deteriorating (e.g. Waycott et al., 2009; Deegan et al., 2012; Alleway and Connell, 2015), information on highly productive habitats is particularly valuable for conservation and management (Seitz et al., 2014; Vasconcelos et al., 2014).

Habitat-specific juvenile survival rates are an important attribute of productive habitats (Beck et al., 2001; Heck Jr, Hays and Orth, 2003; Minello et al., 2003). As survival rates are especially low during early juvenile stages, habitat-specific differences in survival rates can lead to substantial variation in secondary production of juveniles and affect population dynamics at large spatial scales (Lipcius et al., 2019). Hence, quantifying habitat-specific and life-history stage survival rates are vital to properly estimate the function and value of estuarine habitats.

Tethering–a method that restrains a prey to a fixed location for a period of time–is commonly utilized to quantify relative survival rates in field experiments. Tethering allows a researcher to be absent during field trials, thereby reducing the probability of unnatural prey behavior induced by humans (Peterson and Black, 1994; Baker and Waltham, 2020). Tethering in field settings also provides prey with a wider range of movement and the ability to feed and hide as opposed to alternative methods such as caging (Aronson and Heck Jr, 1995; Aronson, Heck and Valentine, 2001). However, tethered animals are more vulnerable to predation due to restricted movement and altered escape behaviors which may artificially reduce survival (Peterson and Black, 1994). Hence, this method can bias survival rates. Consequently, predation rates from tethering are interpreted as relative, not absolute (Aronson, Heck and Valentine, 2001). Moreover, a major assumption that enables comparison of survival across habitats is that the bias associated with the tethering process is consistent across all treatments (Peterson and Black, 1994). Thus, it is imperative that treatment-specific biases such as the interactions between characteristics of the habitats and the tethering procedure be addressed prior to any survival study utilizing tethering (Peterson and Black, 1994; Baker and Waltham, 2020).

The blue crab, *Callinectes sapidus*, is a widely distributed marine and estuarine species. Within Chesapeake Bay, the blue crab is a dominant benthic predator, which may exert top-down control on marine invertebrate communities (Lipcius et al., 2007), as well as a valuable food source for many commercially important species (Stehlik and Meise, 2000; Walter and Austin, 2003; Lipcius and Latour, 2006; Geraldi and Powers, 2011). The blue crab supports one of the most valuable fisheries in the Western Atlantic and Gulf of Mexico; in 2019 US annual commercial landings of blue crab were 66,497 mt valued at US $205.6 million (NOAA, 2019). However, the blue crab population in Chesapeake Bay has fluctuated over the past two decades, reaching an all-time low of 227 million estimated crabs in 2022, the lowest number since the Blue Crab Winter Dredge Survey’s conception in 1990 (Lipcius, 2022).

Blue crabs exploit numerous habitats in their early life stages, such as seagrass beds (e.g. eelgrass *Zostera marina* beds in the Chesapeake Bay), *Spartina alternaflora* salt marshes, and coarse woody debris (Lipcius et al., 2007). Previous studies estimating survival in post-recruitment juveniles have primarily focused on seagrass meadows, algae and unstructured sand habitats to infer nursery status (Pile et al., 1996; Hovel and Lipcius, 2001, 2002; Lipcius et al., 2005; Johnston and Lipcius, 2012; Bromilow and Lipcius, 2017). However, mounting evidence suggests that alternative habitats such as salt marshes serve as highly productive nurseries (Johnson and Eggleston, 2010; Hyman et al., 2022, 2023). While past experiments have not found treatment-specific interactions among unstructured sand and seagrass meadows (Hovel and Lipcius, 2001, 2002; Pile et al., 1996), such interactions have not been quantified in salt marsh nurseries (Shakeri et al., 2020). Given marked structural differences between *S. alterniflora* and *Z. marina* vegetation, assuming that treatment-specific bias is negligible in salt marshes based on tests of treatment-specific bias in seagrass beds is capricious.

In this study, we examined survival of tethered and untethered juvenile blue crabs in artificial salt marsh and unstructured sand habitats with mesocosm experiments. Our objective was to determine whether treatment-specific bias was present among unstructured sand and salt marsh habitat. Moreover, salt marshes are not homogeneous with respect to structural complexity. Thus, we also tested for treatment-specific bias across a range of salt marsh shoot densities observed in the field.

### Logical framework

Under an Information Theoretic framework (Burnham and Anderson, 2002) we developed multiple alternative hypotheses (H_i_) (Chamberlain et al., 1890). Herein we describe and justify the hypotheses and corresponding independent variables.

**H_1_**: Tethering reduces survival, as restraining crabs should increase vulnerability to predation (Pile et al., 1996; Hovel and Lipcius, 2001, 2002).
**H_2_**: Survival increases with marsh shoot density, based on reduced encounter rates or capture efficiency for predators in structurally complex habitats (Shakeri et al., 2020).
**H_3_**: Survival is an additive function of tethering status and shoot density.
**H_4_**: Survival is a function of tethering status, marsh shoot density, as well as the interaction effect of both variables. Juvenile blue crabs may have different escape strategies under differing habitat conditions, such as crypsis in structured habitat vs. escape in unstructured habitat (Peterson and Black, 1994; Zimmer-Faust et al., 1994). Tethering juveniles under different shoot densities may result in non-additive effects on survival (i.e., treatment-specific bias).
**H_5_**: Survival is a function of tethering, shoot density, and predator size, such that survival is inversely related to predator size because larger crabs may be restricted in movement more than small juveniles and as a result less efficient when foraging for small juveniles (Arnold, 1984; Blundon and Kennedy, 1982; Hill and Weissburg, 2013; Shakeri et al., 2020). We further partitioned H1_5_ into two separate hypotheses, one with the effect of predator size within an additive model (H1_5a_) and the other with the effect of predator size plus a tethering-shoot density interaction (i.e., treatment-specific bias; H_5b_).
**H_6_**: Survival is a function of tethering, shoot density, predator size, and prey size. As juvenile blue crabs grow, they are less susceptible to predation as their carapace widens and becomes harder, spines become more prominent, and aggressive behavior intensifies (Hines and Ruiz, 1995; Hovel and Lipcius, 2002; Lipcius et al., 2007; Bromilow, 2017). H_6_ also considered both an additive model (H_6a_) and another with the tethering-shoot density interaction (H_6b_).

## Methods

### Experimental design

The experimental design in this study follows methods employed in Hyman et al. (2023). Eight 160-L recirculating cylindrical fiberglass tanks with a bottom area of 0.36 m^2^, aerated by an airstone, simulated an estuarine marsh. Marsh grass densities were simulated using wooden dowels (1 cm diameter, 30.8 cm height) placed into a 40.6 cm x 40.6 cm plastic pegboard, which was buried 3-5 cm beneath sand from the York River. A control treatment included a peg board without dowels. Juvenile blue crabs were caught weekly using dip-nets in local seagrass and algal habitats. Prior to the experiment, juvenile blue crabs were set into tanks without a predator and recovered 24 h later after draining the tank completely (n = 14). All animals were found within 5 min of searching, which validated the assumption that missing animals at the conclusion of a trial were eaten. Four shoot density treatments with 0, 64, 96, and 128 (corresponding to densities of 0, 388, 582, and 776 m^−2^, respectively) were employed based on densities in *S. alterniflora* salt marshes Cranford, Gordon and Jarvis (1989); Dai and Wiegert (1996); Chaisson, Jones and Warren (2022). In each tank, PVC aqueducts continuously supplied river-sourced water at a constant flow rate. Water level was controlled using a PVC standpipe in the center of each tank (Fig. 1). Tank water was changed completely after each trial (i.e. every 48 h) to reduce buildup of ammonia, nitrates, and other waste compounds. Before each trial, temperature, salinity, and DO were measured in each tank with a YSI data sonde to account for natural fluctuations in the river-sourced flow-through water.

**Fig 1.**
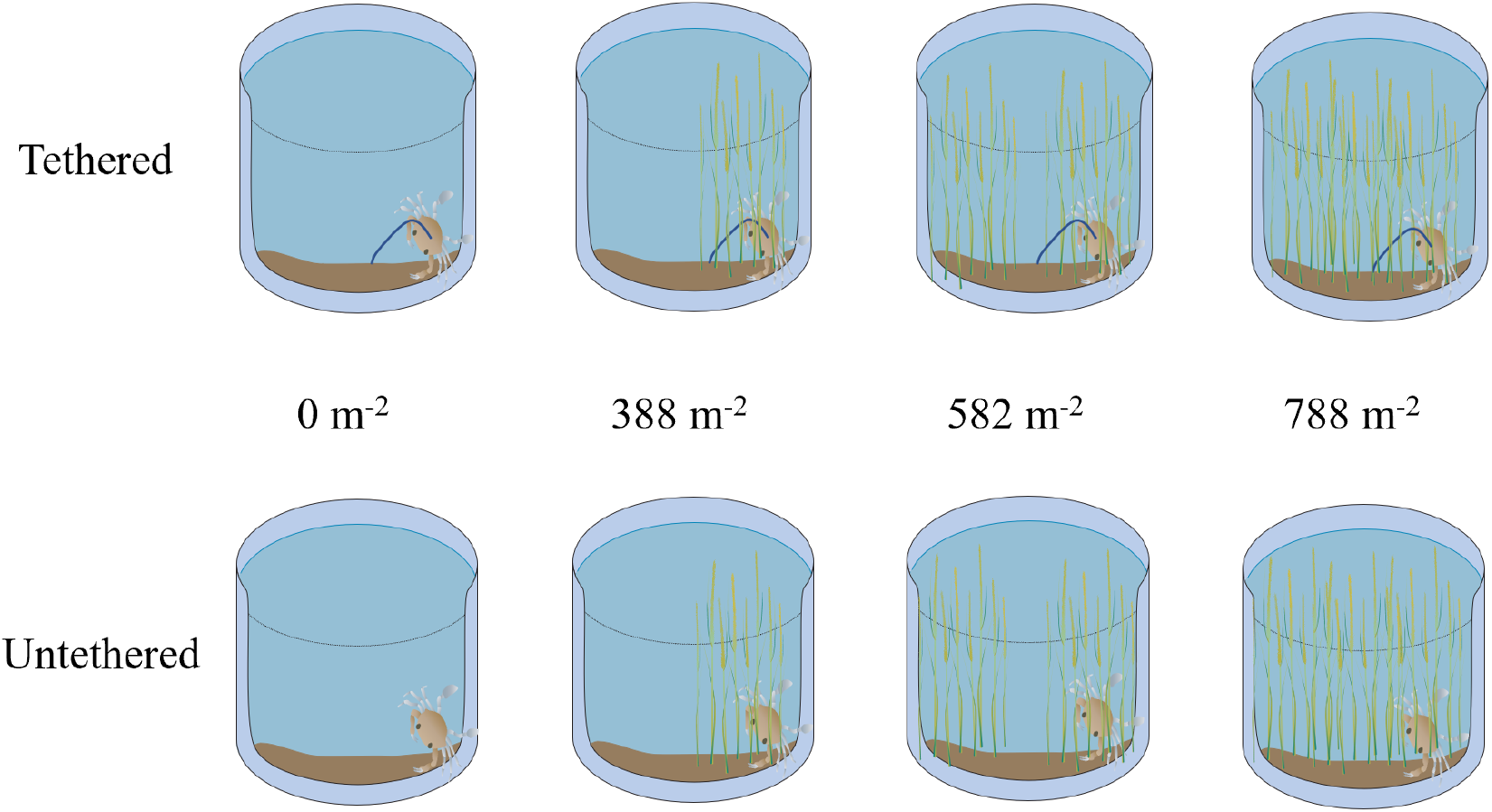
Conceptual diagram depicting mesocosm experimental design and idealized tank setup for tethering-shoot density setup. Both shoot density and tethering treatments were randomized regularly to avoid tank-specific bias.

Adult blue crabs were selected as model predators, as adult conspecifics are among the most important predators of small juveniles (Moody, 2003; Eggleston, Bell and Amavisca, 2005; Hines, 2007; Lipcius et al., 2007; Bromilow, 2017; Longmire et al., 2021). Adult crabs were caught using crab traps in the York River. Following capture, each crab was measured, tagged, placed in a holding tank, and fed juvenile blue crabs to acclimate predators to prey. Holding tanks employed the same flow-through river water as experimental tanks. If an adult crab molted, it was allowed to harden for at least 2 d prior to a trial. Adult crabs were acclimated to experimental conditions, in separate cages to deter antagonistic behavior, for 14 d prior to a trial. Juvenile blue crabs (prey) were acclimated in a separate tank with individual compartments to deter antagonistic behavior.

For each trial, an adult blue crab (predator) was selected randomly from the holding tank and placed into each experimental tank. After recording its carapace width, a juvenile blue crab was placed in the center of each tank near the drainpipe. Each trial ran for 24 h, after which the adult crab was removed with a dip net, placed in a separate tank, fed, and left to acclimate for 24 h before the next trial. Subsequently, each tank was completely drained via a plastic siphon and searched for 10 min (i.e. twice the duration estimated to recover surviving juveniles) for surviving juveniles or carapace fragments. The absence of a juvenile crab or presence of carapace fragments was interpreted as a predation event. Then, tanks were refilled. If an adult crab behaved abnormally, another adult was selected randomly from a holding tank containing replacement crabs.

For the tethering treatments, four juvenile blue crabs were randomly selected 24 h prior to a trial. To adhere the tether, cyanoacrylate glue (super glue) was applied to the dorsal side of the carapace of juveniles between the anterior pair of walking legs to not inhibit movement. A 25 cm thread of monofilament fishing line (4.5 kg test) was set into the glue (Hovel and Lipcius, 2001; Lipcius et al., 2005). A small square of brown duct tape was then applied over the thread and glue to secure the glue-filament base. After tethering, each prey was moved into a holding tank to acclimate before the start of a trial.

At the start of each trail, one juvenile blue crab–either tethered or untethered–was randomly assigned to a tank, such that one tethered and one untethered juvenile crab were in each shoot density replicate (n = 2 tether treatments x 4 shoot density treatments = 8). Water depth was held constant at 24 cm, simulating high tide conditions in local salt marshes. Every two weeks, shoot density treatments were randomized to control for tank effects. This experiment was replicated for 11 trials (n = 8 tanks x 11 trials = 88, Fig. 1).

### Data analyses

After each trial, data were recorded digitally. All data analyses, transformations, and visualizations were completed using the R programming language for statistical computing (R Core Team, 2022). At the conclusion of each experiment, binary survival data were analyzed using generalized linear regression mixed-effects models to evaluate effects of experimental treatments and water chemistry variables. The response variable (probability of juvenile survival) was modeled using a binomial distribution and related to predictor variables using the logit-link (i.e. logistic regression). Predator ID, trial number, and tank ID were initially included as random-intercept effects but discarded due to negligible residual variation explained in all cases, which reduced the model structures to generalized linear models.

The hypotheses for each experiment were translated to sets of statistical models (*g_i_*; Table 2) and evaluated within an Information Theoretic framework (Burnham and Anderson, 2002; Anderson, 2008). Salinity, temperature, and dissolved oxygen (DO) were initially included as fixed effects to ensure that variation in these variables did not influence survival, and were subsequently eliminated from consideration. For each model set, AIC (Akaike’s Information Criterion) corrected for small sample size (AICc) was employed to evaluate the degree of statistical support for each model. Weighted model probabilities (w_*i*_) based on Δ_*i*_ values were used to determine the probability that a particular model was the best-fitting model in a set. Models with Δ_*i*_ values within two points of the best fitting model were considered to have comparable support and further evaluated using likelihood ratio *X*^2^ tests to determine their importance Burnham and Anderson (1998, 2002). When two models had comparable Δ_*i*_ values and likelihood ratio *X*^2^ tests did not suggest significant differences in explanatory power, the simpler model was chosen as the more appropriate model under the principle of parsimony.

## Results

A total of 88 tank-trial combinations were run between June and July 2022, although only 81 were used due to various logistical issues. Data and ranges for physicochemical variables (DO, temperature, and salinity) and sizes of prey and predators are detailed in Table 1 and Figure 2.

**Table 1.** Summary statistics for physicochemical and biological variables Salinity (Sal), Temperature (Temp), Dissolved Oxygen (DO), Predator Width (Predator CW), and Prey Width (Prey CW). CW = carapace width.

| Estimate | Sal | Temp<br>(°C) | DO<br>(mg/L) | Predator CW<br>(mm) | Prey CW<br>(mm) |
| --- | --- | --- | --- | --- | --- |
| Minimum | 16.91 | 23.60 | 5.60 | 104.00 | 11.30 |
| Mean | 19.15 | 24.97 | 6.38 | 132.21 | 20.24 |
| Maximum | 20.35 | 26.20 | 7.75 | 155.00 | 30.20 |

**Fig 2.**
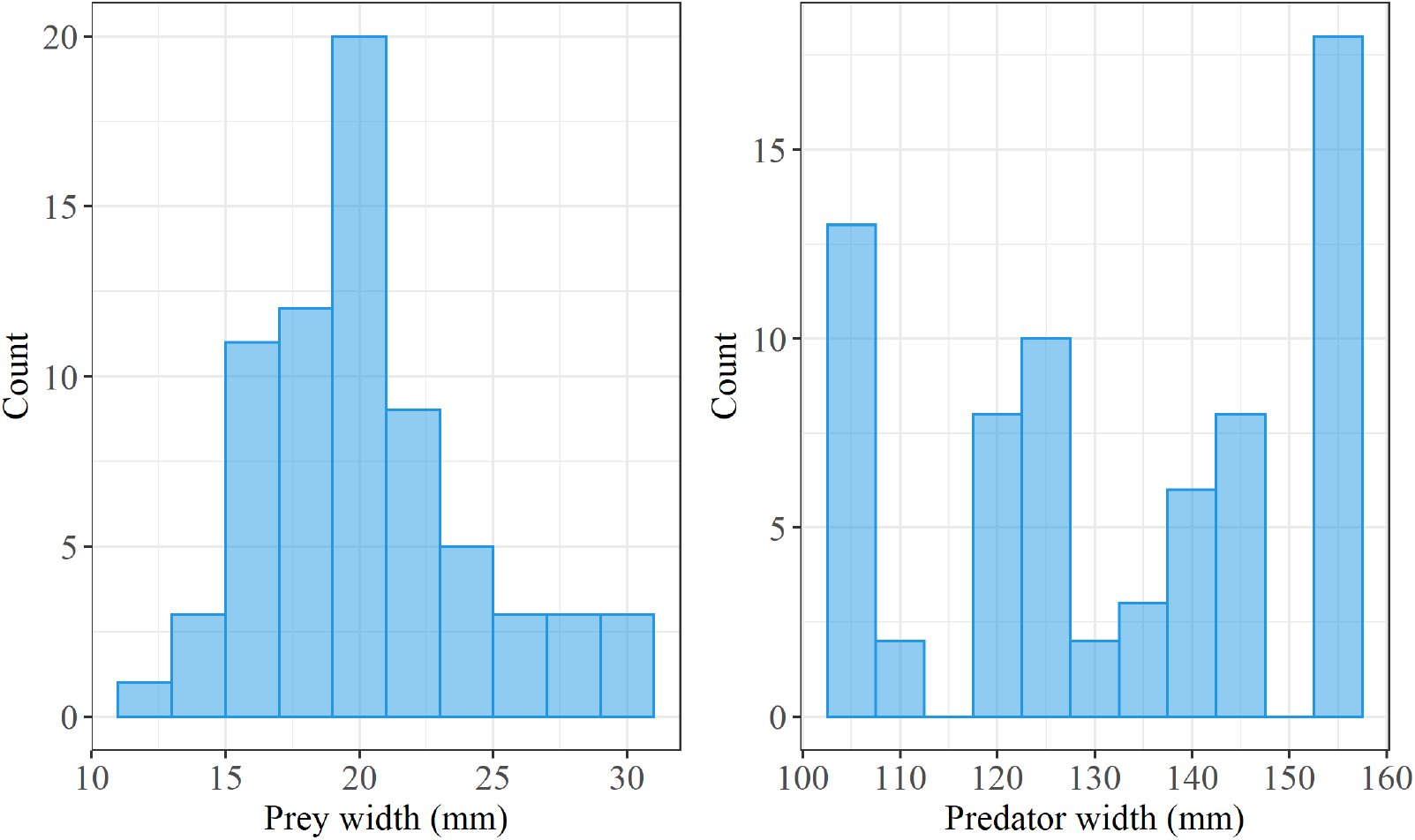
Histogram of all crab carapace widths used in the mesocosm study. Size bins for prey and predator histograms are 2 mm and 5 mm, respectively.

### Model selection

The best model was *g*_3_, an additive model with tethering and shoot density as its predictors (Table 2). All other models except *g*_4_ and *g*_5_ were eliminated from consideration because their weighted probabilities were less than or equal to 0.1. Although models *g*_3_ and *g*_5_ had similar AICc values, model *g*_5_ did not improve predictive performance (likelihood ratio *X*^2^ test, p = 0.16) and the added parameter in model *g*_5_, predator width, was not statistically significant (p = 0.17). Hence, we selected model g_3_ as the most appropriate model.

**Table 2.** Information theoretic analysis of 8 logistic regression models (*g_i_*) using tethering status (T), shoot density (SD), predator size (P), and prey size (Sp) as predictors of juvenile blue crab survival, where AIC_*c*_ is the Akaike information criterion corrected for small sample size, Δ_*i*_ is the difference between any model and the best model in the set, and w_*i*_ is the weighted model probability that a given model is the best among the set considered. Values from the selected model are presented in bold font.

| Hypothesis | Model: Formula | $AIC_c$ | $\Delta_i$ | $w_i$ |
| --- | --- | --- | --- | --- |
| $H_1$ | $g_1$: T | 97.23 | 4.59 | 0.03 |
| $H_2$ | $g_2$: SD | 95.08 | 2.44 | 0.09 |
| $H_3$ | $g_3$: **T + SD** | **92.64** | **0.00** | **0.30** |
| $H_4$ | $g_4$: T + SD + (T x SD) | 94.62 | 1.98 | 0.11 |
| $H_{5a}$ | $g_5$: T + SD + P | 92.93 | 0.29 | 0.26 |
| $H_{5b}$ | $g_6$: T + SD + (T x SD) + P | 95.23 | 2.59 | 0.08 |
| $H_{6a}$ | $g_7$: T + SD + P + Sp | 94.74 | 2.10 | 0.10 |
| $H_{6b}$ | $g_8$: T + SD + (T x SD) + P + Sp | 97.13 | 4.49 | 0.03 |

We also compared *g*_3_ and *g*_4_ – *g*_3_s companion model which included the shoot density-tethering interaction term – to test for treatment-specific bias. Model *g*_4_ with one more parameter did not improve fit to the data (likelihood ratio *X*^2^ test, p = 0.60) and the interaction effect was not statistically significant (p = 0.61). Thus, we conclude that treatment-specific bias was absent.

In model *g*_3_, survival increased with shoot density, such that for every unit (1 shoot) of shoot density the odds of survival increased by 0.23% (Fig. 3). In contrast, tethered crabs were significantly less likely to survive by 111% relative to free-swimming crabs (Table 3).

**Table 3.** Summary results of model *g*_3_ coefficients: T denotes tethering and SD denotes shoot density.

|  | Estimate | Std. Error | z value | P-value |
| --- | --- | --- | --- | --- |
| Intercept | -0.28 | 0.50 | -0.55 | 0.58 |
| SD | 2.22e-3 | 0.9e-3 | 2.48 | 0.01 |
| T | -1.11 | 0.53 | -2.09 | 0.04 |

**Fig 3.**
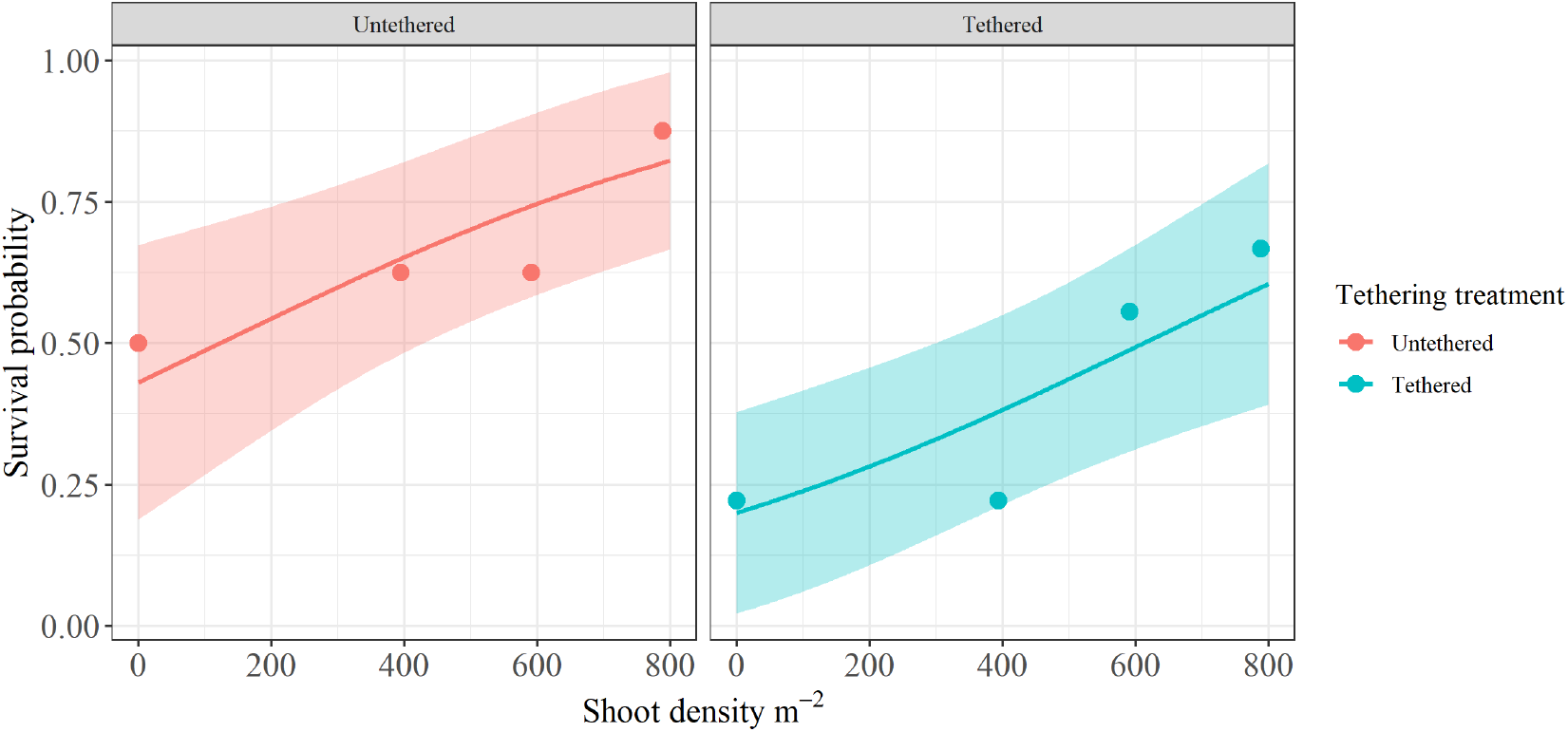
Logistic regression mean conditional effects of shoot density and tethering status (tethered and free-swimming) on survival based on estimates from model g_3_. Points depict aggregated data; mean survival proportions across tethering treatments and shoot densities. Shaded regions denote 95% confidence bands.

## Discussion

This study examined the relationships between juvenile blue crab survival and environmental variables in an artificial salt marsh habitat. The objectives were to test for treatment-specific bias between tethering and salt marsh habitat in mediating survival of juvenile blue crabs under a natural range of marsh grass shoot densities. Our best model was an additive model describing survival as a function of shoot density and tethering status. All variables were informative in explaining variation in juvenile survivorship.

The major findings of this study were that (1) tethering had a significant negative effect on survival, and (2) treatment-specific tethering bias across habitats was not present. The negative effect of tethering was consistent with literature and supports the common assertion that tethering results in salt marsh habitats are a reliable, relative measure of habitat-specific survival (Peterson and Black, 1994; Aronson and Heck Jr, 1995; Aronson, Heck and Valentine, 2001; Hovel and Lipcius, 2002). In addition, there was no significant interaction effect between tethering and shoot density on crab survival, demonstrating that treatment-specific bias was not present irrespective of shoot density. Taken together with earlier studies reporting no treatment-specific tethering bias among seagrass meadows and unstructured sand habitats (Pile et al., 1996; Hovel and Lipcius, 2001), our findings expand the portfolio of habitats under which tethering may be reliably employed with respect to juvenile blue crabs. A benefit of the present study is that results presented here retroactively support past research that did not test for the presence of treatment-specific bias in salt marshes (Hyman et al., 2022; Shakeri et al., 2020).

Survival was positively associated with shoot density, which is consistent with literature (Lipcius et al., 2001; Hovel and Lipcius, 2001, 2002; Hyman et al., 2023) and indicates that variation in structural complexity mediates survival of juvenile blue crabs as for other marine and estuarine juveniles (Hovel and Lipcius, 2002; Long, Sellers and Hines, 2013). In concert with evidence of high abundance and growth of juvenile blue crabs in salt marshes (e.g. Thomas, Zimmerman and Minello, 1990; Fitz and Wiegert, 1991; Zimmerman, Minello and Rozas, 2002; Lipcius et al., 2005; Seitz, Lipcius and Seebo, 2005; Johnson and Eggleston, 2010; Isdell et al., 2021; zu Ermgassen et al., 2021; Hyman et al., 2022), our findings on survival support the conclusion that structurally complex *S. alterniflora* habitats enhance survival rates and promote high abundance of small juvenile blue crabs.

A caveat is that this study used adult blue crabs as the sole predator. Although adult blue crabs are major predators of juvenile blue crabs, it would be worthwhile to validate the findings for piscine predators, which are also major consumers of juvenile blue crabs.

The findings of this study represent important progress in understanding the function and value of alternative blue crab nursery habitats. Seagrass is considered the preferred nursery habitat for juvenile blue crabs (Lipcius et al., 2007). However, the dominant seagrass species of the southern Chesapeake Bay, eelgrass *Z. marina*, is threatened due to increasing temperatures and poor water quality (Orth et al., 2010; Moore, Shields and Parrish, 2014; Patrick et al., 2018), and may be eliminated in the near future (Waycott et al., 2009; Wilson and Lotze, 2019). Mounting evidence suggests salt marshes and certain unstructured sand habitats may be as valuable or more so at the population level due to their extensive areal cover (Lipcius et al., 2005; Johnson and Eggleston, 2010; Hyman et al., 2022). Hence, it is critical that alternative nursery habitats, such as salt marshes, are protected for their contribution to blue crab populations.

## Acknowledgments

The authors thank Michael Seebo for mesocosm setup and advice on capturing, acclimating, and maintaining experimental animals. Authors also acknowledge the Virginia Institute of Marine Science’s NSF Research Experience for Undergraduates program.

